# Bayesian Multivariate Longitudinal Model for Immune Responses to Leishmania: a tick-borne Co-Infection Study

**DOI:** 10.1101/2022.06.20.496877

**Authors:** Felix M. Pabon-Rodriguez, Grant D. Brown, Breanna M. Scorza, Christine A. Petersen

## Abstract

While many Bayesian state-space models for infectious disease processes focus on population infection dynamics (e.g., compartmental models), in this work we examine the evolution of infection processes and the complexities of the immune responses within the host using these techniques. We present a joint Bayesian state-space model to better understand how the immune system contributes to the control of *Leishmania infantum* infections over the disease course. We use longitudinal molecular diagnostic and clinical data of a cohort of dogs to describe population progression rates and present evidence for important drivers of clinical disease. Among these results, we find evidence for the importance of co-infection in disease progression. We also show that as dogs progress through the infection, parasite load is influenced by their age, ectoparasiticide treatment status, and serology. Furthermore, we present evidence that pathogen load information from an earlier point in time influences its future value and that the size of this effect varies depending on the clinical stage of the dog. In addition to characterizing the processes driving disease progression, we predict individual and aggregate patterns of Canine Leishmaniasis progression. Both our findings and the application to individual-level predictions are of direct clinical relevance, presenting possible opportunities for application in veterinary practice and motivating lines of additional investigation to better understand and predict disease progression. Finally, as an important zoonotic human pathogen, these results may support future efforts to prevent and treat human Leishmaniosis.

**AUTHOR SUMMARY:** The immune system is a complex network that involves organs, cells, and proteins working together with the main purpose of protecting the body against harmful microorganisms such as bacteria, viruses, fungi, and toxins. To explore and study the responses of the host immune system during the course of a disease, we modeled the interaction between pathogen load, antibody responses, and the clinical presentation of this complex system. Specifically, we focused on *Canine Leishmaniasis* (CanL), a vector-borne disease caused by a parasite that affects internal organs of the body and is known to be fatal if patients remain untreated. In addition, we also considered the impact of possible co-infections with other diseases, which could potentially interact with many disease processes and contribute to different outcomes for infected subjects. With CanL specifically, we consider the presence of *Borrelia, Anaplasma, Ehrlichia*,, and Heartworm. In general, one limitation in vaccination strategies is a focus on neutralizing antibodies, without incorporating the broader complexities of immune responses. Here, we explore this complexity by jointly considering the interaction between pathogen and antibody development with the purpose of improving our understanding of the processes of disease progression and natural immunity.

In this paper, we present a Bayesian model specification for immune responses to a *Leishmania* infection considering a tick-borne co-infection study. The model implementation is based on the general vector autoregressive (VAR) model, adapted to the problem under study. While the methodology around Bayesian VAR models is not new in the literature, in this work we adapt the more general VAR approach in a parsimonious way to a particular subclass of longitudinal problems. We believe our defined Bayesian model is useful to clinicians and veterinarians to better understand the immune responses and *Leishmania* infection control over time, which makes this work a novel application of Bayesian VAR models. We present evidence that pathogen load information from an earlier point in time influences its future value and that the size of this effect varies depending on the CanL clinical stage of the dog. In addition to characterizing evidence for the processes driving disease progression, we predict individual and aggregate patterns of CanL progression.

The structure of this paper starts in Section 2 with an introduction to CanL infection as well as a discussion of possible co-infection with other pathogens. In Section 3, we include a description of the motivating prospective study along with the measured individual-level variables, a definition of the clinical signs of leishmaniosis infection, and a description of the available data coming from the study. In addition, this section explains the dynamic process and corresponding model specification via Bayesian methodology and a statement of contribution. A summary of prior distributions for model parameters, model implementation details, and convergence diagnostics are also included. In Section 4, we provide summary results from the posterior distribution as well as a summary of the corresponding disease progression forecasts. In Section 5, we discuss the results and describe future considerations to improve and extend the model.

## 2 INTRODUCTION: CANINE LEISHMANIASIS

Leishmaniasis is a life-threatening parasitic disease endemic in 98 countries, with an estimated 50,000 - 90,000 new cases annually, according to the World Health Organization (WHO). Despite this widespread distribution, *Leishmania* infection is considered a neglected tropical disease. There is no approved human vaccine, and current treatments can have adverse side effects and do not lead to a clinical cure. This can lead to the recrudescence of disease^1^. *Leishmania* infection is caused by a family of parasites in the *Leishmania (L*.*) donovani* complex, including *L. infantum*. It is known that *L. infantum* is a zoonotic infectious disease with dogs as the predominant reservoir. Spatial modeling has shown that human Leishmaniasis incidence positively associates with the prevalence of *L. infantum* infection among dogs in an endemic area^2^. Therefore, limiting transmission among and from infected dogs is an important target for preventing human infections. However, our understanding of CanL transmission dynamics is hampered by a lack of ability to accurately predict clinical CanL progression within the canine reservoir.

In addition to helping understand *L. infantum* transmission, elucidating CanL clinical progression dynamics can give us important pathophysiological insight into human *Leishmania* infection, as the canine immune response to *L. infantum* closely mirrors human disease^3^. In both humans and dogs, the majority of *L. infantum* infections are sub-clinical, controlled by a Type 1 immune response. However, dogs can still be infectious to the insect vector during this time^4,5^. For reasons that are not well defined, clinical progression occurs in a subset of dogs and people, where Type 1 immunity wanes and parasite load increases. Such progression leads to increased transmission potential.

Diagnosis of CanL is based on a combination of molecular and clinical factors. Molecular diagnostics include serology, which may indicate current infection or previous exposure, and Polymerase chain reaction (PCR) of *Leishmania* deoxyribonucleic acid (DNA) from whole blood, which indicates current infection above the threshold of detection. Increasing parasitemia level is associated with disease progression^6^. Anti-*Leishmania* antibodies produced during infection are generally non-protective, and increasing serological titers are also associated with worsening disease status^7^. As infected dogs progress to early CanL, clinical signs of disease are non-specific (i.e. weight loss, lymphadenopathy, lethargy) and molecular diagnostics may or may not be positive. As the disease progresses, anemia and hypergammaglobulinemia usually precede chronic kidney disease and ultimately kidney failure if untreated^7^.

In this work, we prospectively followed a cohort of dogs with sub-clinical *L. infantum* infection and collected longitudinal molecular diagnostic and clinical data to evaluate disease progression over 18 months. We used this data to develop a joint Bayesian state-space model to predict progression kinetics based on continuous diagnostic data and discrete clinical stages. We explore population progression rates and present evidence for important dependencies among measured components of disease status. In particular, we find further evidence for the importance of co-infection status in CanL progression. Furthermore, we show that as dogs advance through the infection, their age, ectoparasiticide treatment indicator, and serology all impact parasite load. We also show that pathogen load information from a previous time influences its future value and that the magnitude of this effect varies depending on the dog’s clinical state, where the effects of these clinical drivers are being measured via the model parameters. We also look into the model’s suitability as a tool for predicting clinical disease progression.

One of the primary non-time-varying predictors of interest within the study is the output of the IDEXX 4Dx Plus SNAP test, which is a combined in vitro test for the detection of antigen to *Dirofilaria immitis*, antibodies to *Borrelia burgdorferi, Anaplasma phagocytophilum, Anaplasma platys, Ehrlichia canis*, and *Ehrlichia ewingii* in canine serum, plasma, or anticoagulated whole blood. It is known that previous or simultaneous infections of a host by multiple pathogen species (*spp*.) can complicate CanL clinical outcomes^8^. Some studies have shown that the presence of additional interacting pathogens can modify the immune response of the affected hosts^9^. This phenomenon of co-infection is of great importance in the progression of CanL^10^. By studying the effect of this predictor, we aim to measure the strength of its effect during the course of infection and identify different possible clinical outcomes for co-infected subjects. Within our CanL cohort, we specifically considered the effects of *Borrelia burgdorferi, Anaplasma spp*., *Ehrlichia spp*., and Heartworm exposure.

## 3 MATERIALS AND METHODS

### 3.1 Cohort Selection and Diagnostics

A prospective study was performed using a cohort of naturally *L. infantum* exposed, client-owned hunting dogs from the United States. This cohort was selected in such a way that canine subjects have not yet progressed to severe disease and were not recently exposed to selected tick-borne co-infections. Specifically, the inclusion criteria included dogs that must be IDEXX 4Dx Plus SNAP test seronegative, indicating no recent exposure to common tick-borne pathogens: *Borrelia burgdorferi, Anaplasma spp*., or *Ehrlichia spp*. To be included, dogs also had to be positive on one Leishmania diagnostic test: Real-Time quantitative PCR (RT-qPCR) of whole blood or Dual Path Platform ® (DPP) Canine Visceral Leishmaniasis serological test^11,12^, or have a Leishmania diagnostic positive dam or full sibling, indicative of infection or in utero exposure to *L. infantum* respectively. On the other hand, exclusion criteria included dogs having more than 2 clinical signs of CanL or DPP reader value greater than 200, indicating clinical manifestation or progressive disease. Dogs that were seropositive on the IDEXX 4Dx Plus SNAP test were also excluded, indicating recent or concurrent exposure to a tick-borne pathogen. A total of fifty dogs were enrolled in the study and followed over the course of 18 months. The cohort was equally randomized into a tick prevention ectoparasiticide treatment or placebo group under a double-blind setting, where the age range of dogs was from 0 to 11 years old. Using these criteria, we established a cohort, allowing us to monitor the progression from no clinical manifestation to showing clinical signs of canine leishmaniosis in the presence or absence of nascent tick-borne co-infections.

### 3.2 Variables

During the data collection period, diagnostic variables and clinical disease were assessed at three-month intervals. This included RT-qPCR for parasite quantification in the blood of subjects. Pathogen load is then presented as the number of parasite equivalents per mL of blood, calculated from a standard curve of canine blood spiked with a known number of *L. infantum* promastigotes^11^. Relative quantification of anti-*Leishmania* antibody levels in subject sera were performed using an indirect Enzyme-linked immunoassay (ELISA) against soluble *Leishmania* antigen (SLA). Wells of ELISA plates were coated with 200 ng/mL SLA obtained by repeated freeze-thaw of *L. infantum* promastigotes and probed with canine serum samples diluted 1:500. The optical density (OD) measured at 450 nm on a plate reader is shown as a ratio to a cut-off OD (average of the control sample ODs + 3 standard deviations). OD ratios *>*1 indicate positive serology. Complete blood count and serum chemistry panels were performed by IDEXX Reference Labs.

The SNAP variable presents dichotomous results from the IDEXX 4Dx Plus SNAP test, which tests for serologic reactivity to tick-borne bacterial antigen from *Borrelia burgdorferi, Anaplasma spp*., and *Ehrlichia spp*., indicating previous exposure to these pathogens. Although all dogs were negative by the 4Dx Plus SNAP test at enrollment, we observed that 44% of the dogs tested positive on the 4Dx Plus SNAP test at some point during the study period, which indicates co-infection between *L. infantum* and a tick-borne pathogen occurred during the study period. While this variable is originally collected at each time point, an overall variable is implemented within the model as a fixed effect. The results from the SNAP test were used to define the dichotomous variable for co-infection status, which assigns a value of 1 if a dog showed signs of co-infection with another pathogen on at least one-time point, or assigns the value 0 if the dog never showed signs of co-infection. In this work, we also considered results from the DPP serological test, which detects anti-*Leishmania* antibodies. DPP is similar to SLA ELISA, but more specifically as it detects antibodies specific for recombinant chimeric k28 protein, and results are measured with a digital intensity reader. The mean and standard deviation in DPP results are 25.20 and 41.56, respectively. However, the DPP test was only performed at enrollment, while the SLA ELISA was performed at three-month intervals. Finally, treatment with the ectoparasiticide medication sarolaner (Zoetis Inc.) or placebo was included. We have that 50% (*n* = 25) of the cohort was randomly assigned to each treatment group at enrollment.

### 3.3 Bayesian Vector Autoregressive models

Vector autoregressive models are used to study the relationships between multiple time series. The main characteristic of these models is the inclusion of lag values of outcome variables as model predictors. That is, for instance, the current disease status of a subject would be modeled using the disease status at previous time points. Reliable estimation of the model parameters can be challenging, especially with small datasets^13^, which is the case of the current study. Bayesian VAR models use prior information about model parameters^14^ to help with the stabilization of the parameter estimation, a task that is sometimes difficult for classical VAR models. The idea is that prior knowledge introduces shrinkage for model parameters. In addition, Bayesian VAR models are also extremely powerful forecasting tools that allow the researcher to place more weight on the information that is provided by the lags of a particular outcome variable.

The work of Koop and Korobilis^15^ provides an introduction to Bayesian multivariate time series methods for the case of macroeconomics, but the usage of this methodology is certainly applicable for a wider range of research areas such as infectious diseases. They present examples of different VAR model implementations as well as time-varying parameter extensions. In addition to simple VAR models, they present several examples of how Bayesian inference requires the use of Markov chain Monte Carlo (MCMC) methods for the development of state space models, by describing the algorithms. Since Bayesian forecasting is a natural product of a Bayesian approach to inference, the work of Geweke and Whiteman^16^ provides a good introduction to the principles of Bayesian forecasting and a description of the advances in computational capabilities.

A more closely related work to the study and prediction of infectious diseases is the work by Seedorf et. al.^17^, which proposed a multivariate Bayesian hierarchical model with autoregressive errors to jointly analyze continuous and ordinal longitudinal data. Interestingly, they also use a longitudinal study of canine leishmaniosis, but their particular attention in the clinical setting is when ordinal outcomes are measured alongside other variable types that may aid clinical decision-making. Their proposed work is particularly applicable when multiple, imperfect measures of disease progression are available within the dataset. The authors’ simulation study shows that when an autoregressive component is present in the data generation process, its inclusion in the model can improve short-term forecasts of both the continuous and ordinal outcomes.

#### 3.3.1 Statement of Contribution

While Bayesian VAR models are well established, we have found that such techniques can be both overparameterized and more difficult to interpret in small sample settings. A more parsimonious approach allows us to extract biologically meaningful parameters, while also reducing the dimension of the parameter space. While what we have proposed does not extend the very general class of VAR models, we feel that it provides a helpful illustration of how such models can be adapted to individual problems. This adaptation of more general VAR models to a particular subclass of longitudinal problems is expected to be useful for modelers collaborating with clinicians and subject matter experts. The novelty of this work comes with a special application to the modeling and forecasting of within-host dynamics of infectious diseases, immunological responses, and disease classification.

### 3.4 Model Specification

The clinical signs of *Leishmania* infection are known to vary widely as a consequence of several pathogenic mechanisms. Depending on the subject, different organs can be affected, and there is a diverse range of immune responses built by individual hosts. This variability and non-specificity of clinical manifestations make diagnosis and treatment challenging. The diagnosis of CanL is performed based on clinicopathological manifestations and by confirmation of infection by using mainly serological and additional molecular techniques. In response to these challenges, Solano-Gallego et al.^7^ proposed a system identified as the LeishVet score, that uses serological results, clinical signs, and laboratory findings to differentiate and categorize patients with CanL into stages. Stage 1 is used for a case of mild disease, Stage 2 for moderate disease, Stage 3 for severe disease, and Stage 4 to identify very severe, end-stage disease. LeishVet is a group of experts that proposed guidelines and recommendations designed primarily to help the veterinary clinician in the management of canine leishmaniosis. This LeishVet score system is helpful because it assists clinicians and veterinarians in determining the appropriate therapy, predicting prognosis, and implementing adequate follow-up steps required to help patients with CanL. In this study, physical exam findings, complete blood count, and serum chemistry values were used to stage CanL severity based on the LeishVet staging guidelines proposed by Solano-Gallego et al.^7^. We have modified the staging system slightly to include a Stage 0 representing no signs of disease.

When it comes to CanL studies, determining the course of treatments is not an easy task, particularly when many unknown relationships between *Leishmania* infection and the immune system still remain unanswered. While various laboratory techniques are beneficial for answering questions pertaining to a particular disease, it is always critical to have in mind that qPCR data should not be separated from data collected from other sources, such as clinicopathological and serological examinations^7,18^, for determining the clinical decision. Thus, rather than focusing on infectious agent transmission between individuals, the Bayesian model presented in this work is thought to be dynamic and interactive within the host.

To assess the CanL disease progression of subjects over time, we took into account the LeishVet stages, pathogen load, and level of anti-*Leishmania* antibodies as explained in the previous section. Each dog was classified based on the scoring proposed by Solano-Gallego et al.^7^. We further aggregate this scoring into the qualitative categories described in Equation 1, for the *i*th dog at time *t* + 1, where we defined disease status *D*_*i*,*t*+1_ for the corresponding indexes. This categorization was encoded as separate indicator variables for each stage over time for each dog.

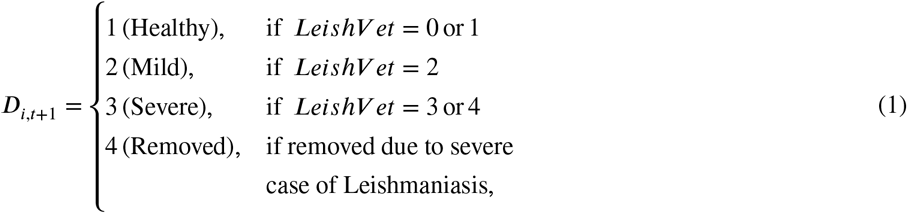

Once an individual is infected with the parasite *L. infantum*, a replication process takes place. At the same time, the host immune mounts a response against the parasite. Therefore, parasitemia may fluctuate as these opposing processes occur, but systematic variation over time may indicate the future trajectory of the disease. One way to measure the pathogen status component of the overall dynamic process is through pathogen burden. For *Leishmania*, let us define *P*_*i*,*t*+1_ to denote the pathogen load for the *i*th dog at time *t* + 1, measured as the number of parasites per mL of blood. Further, let *A*_*i*,*t*+1_ denote the anti-*Leishmania* antibody level measured by ELISA SLA OD ratio for the same indexes. As indicated in the previous section, the disease status was assessed by the proposed variable *D*_*i*,*t*+1_ as defined in Equation 1.

In Figure 1, we illustrate the model’s temporal dependence structure with time index *t* and subject index *i*, which is constructed sequentially and dynamically. This diagram shows how each model component can either directly or indirectly influence each proximal state component. To capture the evolution of state over time, we assume that the future state for each model component is dependent on the full current state of the host (pathogen load, antibody levels, and disease state progression) via some functions, denoted by *f*_*D*_, *f*_*A*_, and *f*_*P*_, describing the expected state at the next time point for each of the three main model components. In addition, the case history or any other fixed effects for the host was also considered to affect the future state, which is encoded by *X*_*i*_, a subject-specific row-vector containing non-time-varying predictors of progression characteristics. This structure could be considered a generalized vector-autoregressive model, though it has been tailored to the expected dependence structure of CanL progression. In general, we have a function *f*_*D*_ describing the disease progression in terms of clinical signs as depending on the current immunopathogenic state, while the functions *f*_*A*_ and *f*_*P*_ describe the antibody levels and pathogen load also as functions of the previous or current state. In addition, these components can incorporate individual-specific error components, capturing measurement error and individual-level heterogeneity, which could be caused by clinical differences associated with the disease. Then, in general, the future states of these components are determined by these update functions, random errors, and parameters associated with the dynamic system, all of this under some distributional assumptions.

**FIGURE 1.**
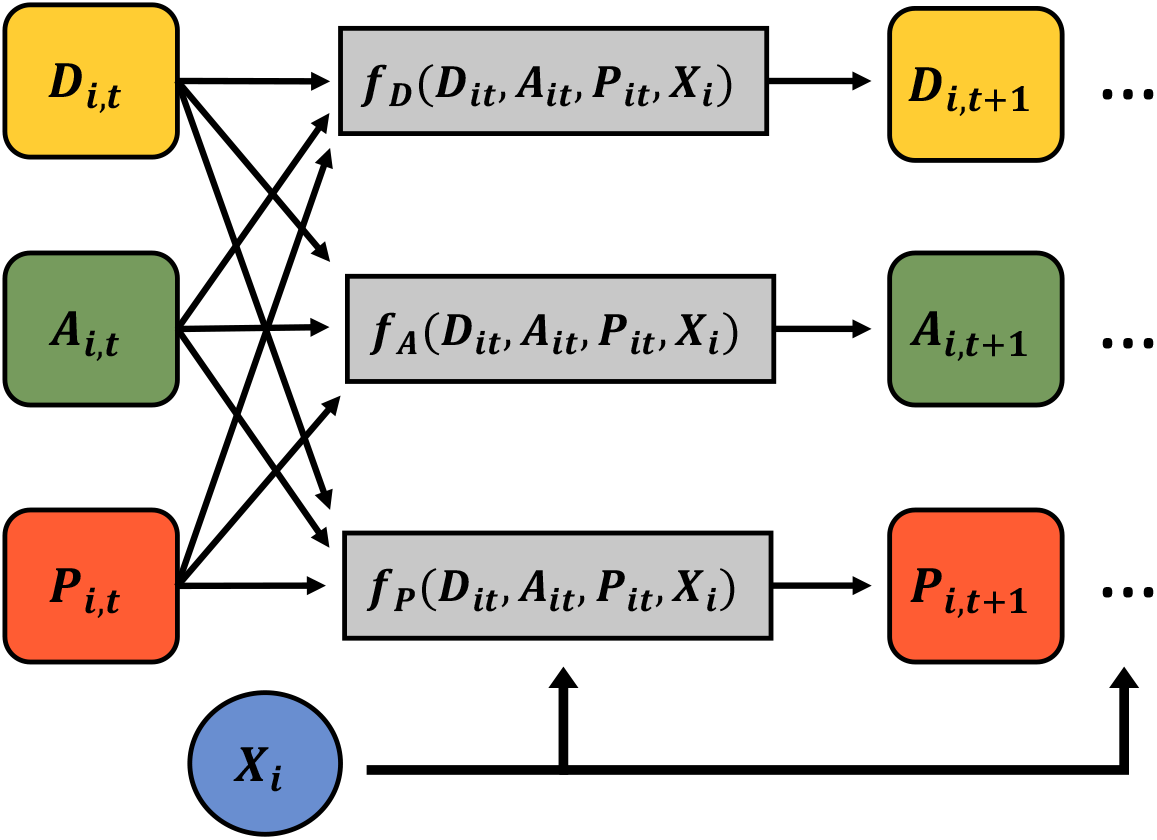
Graphical display of how the disease state (*D*), immune responses (*A*), and pathogen load (*P*) components are related to predicting the future value of each state.

The precise specification of the update functions shown in Figure 1 could be as simple as linear functions of the three state components, in which case the model would reduce exactly to a VAR process. In addition, expansions could be considered (e.g., interactions, non-linear terms, basis expansions, or non-linear link functions coupled to the distribution of subsequent components). We present a simple formulation with Gaussian distributions for pathogen load and antibody response, and a multinomial-logit link for disease status, where functional forms or update functions *f*_*D*_, *f*_*A*_, and *f*_*P*_ are defined as shown in Equation 2.

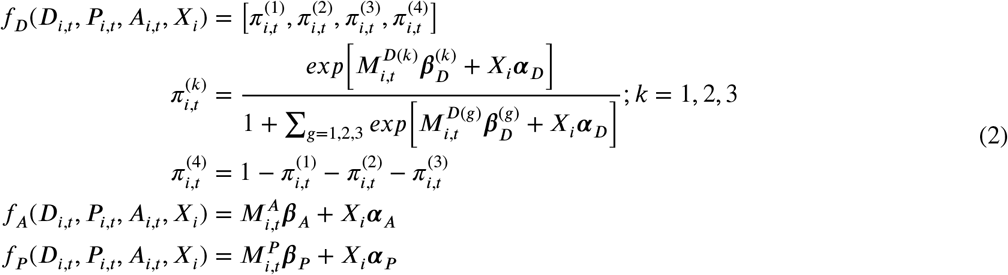

For *N* = 50 dogs and *T* = 7 time points, we have then a conditional layout on dependence structures, where each model component is distributed independently according to Equation 3. In the multinomial distribution, 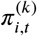 defines the probability that the *i*th dog at time *t* is classified with disease status *k* = 1, 2, 3. The primary components of the model, which are denoted as *P*, *A*, and *D*, have design row vectors that change over study time, denoted by 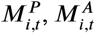, and 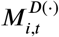, respectively. Non-time-varying predictors are captured in a subject-specific row vector *X*_*i*_ (intercept, age, SNAP, DPP, and an indicator for treatment).

For the evolution of pathogen load, 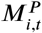 captures the current pathogen load, an interaction between disease stage and pathogen load, an interaction between disease stage and prior changes in pathogen load, and an interaction between disease stage and prior changes in antibody levels. On the other hand, the antibodies are modeled as dependent on current pathogen load and antibody levels as well as in the previously indicated interaction terms. In order to interpret whether a host response to an infection seems appropriate at a particular time point, it needs to be assessed in relation to the pathogen load that triggered the response. By using the current value of pathogen load and its interaction with the disease stage, we are able to understand one of the most fundamental questions, which is the relationship between pathogen load and the severity of the disease. We explore this relationship in more detail by including interaction terms and prior information.

The main difference between the two row vectors 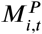 and 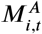 is the assumption that the production of antibodies depends on the production of pathogen load and the current value of antibodies, but the pathogen load is not directly dependent on antibodies, which are generally non-neutralizing. For the qualitative disease stage, 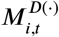 considers the current disease stage and the interaction of the disease stage with the current pathogen load, and the interaction of the disease stage with the current antibody levels. For instance, 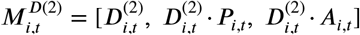. For this model specification, notice that the update functions from Figure 1 are defined to be the mean structures of the distribution.

The dimensions of the different subject-specific row vectors, denoted by *M*_*i*,*t*_, vary by model components. For example, 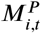 is a 1 × 10 matrix (or row vector), while 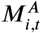 and 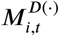 are a 1 × 8 and 1 × 3 row vectors, respectively. Each entry within these row vectors is a scalar, representing different possible drivers of the model dependence. In the case of the subject-specific row vector *X*_*i*_, which includes non-time-varying predictors and an intercept, its dimension is 1 × 5, which includes age group, dichotomous results from SNAP test, DPP results, and an indicator for the treatment group. The elements in *X*_*i*_ are also scalars.

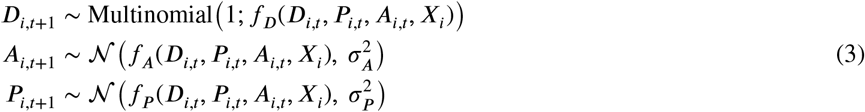

As disease progression for *Leishmaniasis* depends on both pathogen and immunological behavior^7,19^, we included in the row vectors, both components as well as interaction terms with the disease status. To measure the strength of the effects of changing pathogen burden and antibody levels, we computed lag-1 differences for these quantities (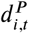 and 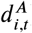). These differences in pathogen load (or antibody levels) are calculated as the difference between readouts at the time *t* and *t* − 1. The row vectors also include the interaction of these lag-1 differences with the clinical stage. Shang et al. showed^20^ that age and external clinical status of dogs are associated with the prevalence of *Leishmania* infection. In their study, dogs that were +1 years old had a higher prevalence of *L. infantum* infection than younger dogs (≤ 1 year old), resulting from most likely longer exposure to infective sand fly bites. We, therefore, included age group and clinical information (e.g. SNAP and DPP results at baseline, treatment group) in our row vector *X*_*i*_. Notice that *X*_*i*_ is included in the three distributions in Equation 3. Finally, the components of the row vector 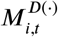 will contribute to the probability that a dog will be classified into a particular disease status. This term is important for measuring the transitions between the four different qualitative categories as described in Equation 1. Overall, this specification is a special case of the well-studied VAR model, which has been applied in fields as diverse as immunology and econometrics^21,22,23^. In this case, we augment the model with a latent component for the disease stage and carefully constrain the temporal dependence structure to encode structural information about the problem, such as the varying effects of shifting antibody levels by disease stage.

Using the model definition as described by Equations 2 and 3, the joint likelihood can be defined by the product of each probability density (or probability mass) function corresponding to each model component. In this case, we have two continuous outcomes (pathogen load and antibody levels) and one categorical (disease status). To facilitate the definition of the likelihood presented in Equation 4, let us define 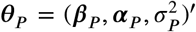 as the set of vectors associated with pathogen load. Similarly, we can define the set of parameters associated with the antibody levels and disease status as 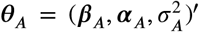 and 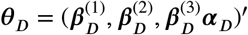, respectively.

For time points *t* = 2, …, *T*, the conditional distributions for pathogen load (*P*) and antibody levels (*A*), denoted by *f* (*P* _*i*,*t*_│·) and *f* (*A* _*i*,*t*_│·) respectively, where *i* indicates subject, depend on autoregressive values (previous information) of the outcomes and their associated parameters ***θ***_*P*_ and ***θ***_*A*_. We assume that these two continuous outcomes follow normal distributions as previously shown in Equation 3, where the mean structures are defined as in Equation 2. The conditional distributions for disease status (*D*), denoted by *f* (*D* _*i*,*t*_│·), follow a multinomial distribution with a logit link function and associated parameters ***θ***_*D*_. In addition, the terms *f* (*P*_*i*,1_), *f* (*A*_*i*,1_), and *f* (*D*_*i*,1_) denote the probability density distributions of the model components at the initial time point (*t* = 1).

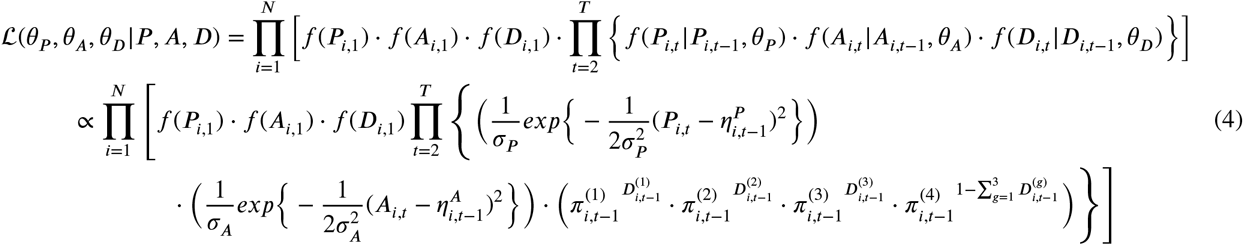

In the likelihood, we have that the mean expressions for each normal density distribution are defined as 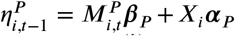, and 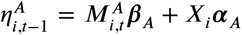, respectively. The expressions for the 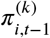 are presented in Equation 2, and where 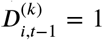 for *k* = 1, 2, 3 if *D*_*i*,*t*−1_ = *k* and 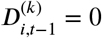 otherwise, which are defined based in Equation 1. Note that a proportionality notation is used in the second line of the likelihood, which helps in factoring out all of the constants or fixed quantities. In addition, notice that we defined our baseline category in the multinomial distribution as 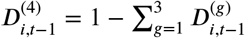

### 3.5 Computation and Model Diagnostics

To fit the Bayesian model presented in this work, we coded and implemented a Metropolis-Hastings within Gibbs algorithm in R^24,25^, to obtain posterior samples of model parameters and latent quantities. A general description of this MCMC approach is provided by Liu^26^ and Carlin^27^. We employed proper prior distributions to impose penalties for large regression weights in order to promote model stability and computational tractability. We used standard independent normal prior distributions, 𝒩,[(0, 1), for the conditional and unconditional effects (***β***’s) as well as for the parameters associated with non-time-varying predictors (***α***’s), in order to shrink these effects towards zero. A gamma prior distribution, Г(1, 1), was used for variance terms (***σ***^2^’s). During the fitting procedure, a scaling approach was used in the continuous longitudinal outcomes and covariates to avoid computational overflow in R due to large values in the readouts. However, results and predictions are provided in the original scale of the corresponding variables. For the case of unobserved or missing longitudinal outcomes (latent quantities), we used Bayesian imputation within the analysis to obtain the posterior distribution of these quantities of interest. This Bayesian approach assumes that the missing data are missing at random (MAR), which means that the probability of missingness depends only on observed data, and not on the missing values themselves after conditioning on the observed data.

Posterior results are based on a set of 3 chains of 25, 000 iterations each where 5, 000 iterations were discarded as burn-in, for a total of 20, 000 iterations on each chain. We ran the MCMC sampler on a laptop computer with an Intel Core i7 processor and 12 GB of memory in order to evaluate the computing performance of our proposed Bayesian approach. We achieved a running time of around 18 hours for the number of iterations per chain ran in parallel, which is acceptable for the complexity of the presented model. While we are aware of the length of the running time, we believe there are numerous opportunities for optimization. For instance, we could consider a different MCMC algorithm such as Hamiltonian Monte Carlo (HMC) or even programming the MCMC sampler into a high-level, high-performance dynamic programming language such as Julia^28^.

To summarize the parameters’ posterior distributions, a table is constructed and presented in Section 4. This table includes the calculation of the posterior means, standard deviation (SD), 95% credible intervals (Cr-I), all based on the 3 chains of the MCMC samples. The limits of the 95% Cr-I are determined in such a way that the lower value is the 0.025 quantile (or the 2.5% percentile) of the posterior distribution, while the upper limit is the 0.975 quantile (or the 97.5% percentile) of the posterior distribution. In addition to these summaries, we also include the posterior probabilities of each parameter (*x*) being either positive or negative, which is *P* (*x >* 0) or *P* (*x <* 0). This is the main criterion used for assessing the strength of evidence for corresponding parameters. To determine whether the effect is moderate or strong, we use the following rule: We consider any parameter with a posterior probability of being positive (or negative) in the interval [0.65, 0.85) to be a moderate effect, while any parameter with a posterior probability of being positive (or negative) in the interval [0.85, 1] will be considered as showing a strong effect. Using this approach, we determine which parameters drive disease progression among the cohort and its dependence on future occasions.

To assess the convergence of the parameters, a Gelman-Rubin diagnostic was computed using the coda package in R^29^. The Gelman-Rubin criterion evaluates convergence by comparing within and between chain variability, which requires at least two MCMC chains to be calculated. The works of Gelman and Rubin^30^ and Brooks and Gelman^31^ suggest that diagnostic values greater than 1.2 for any of the model parameters should indicate a level of non-convergence. However, in practice, a more stringent criterion is often used to declare convergence, where the value of 1.1 is used. Thus, as a rule, to assess convergence in this work, we used the value of 1.1. Therefore, we say that a factor of 1.1 or below indicates that the parameter has reached convergence, which was achieved by all parameters in the model.

We also computed the Monte Carlo Standard Error (MCSE), which is a measure of the precision of the posterior distribution obtained from an MCMC algorithm. The purpose of MCSE is to provide a measure of the accuracy of the estimate of the posterior distribution, which is the distribution of the parameters of interest after taking into account the data and prior information^32^. MCSE is calculated by estimating the standard deviation of the MCMC samples of the posterior distribution, which provides a measure of the variability in the posterior estimates due to the Monte Carlo sampling process. This value is then divided by the square root of the effective sample size (ESS), which is the number of independent samples that the MCMC algorithm generates. This result is a standardized measure of the variability in the posterior estimates, which is expressed in the same units as the parameter being estimated. If the MCSE is small relative to the posterior distribution, this indicates that the estimate is relatively precise, meaning that it is likely to be close to the true value of the parameter. Conversely, if the MCSE is large relative to the posterior distribution, this indicates that the estimate is relatively imprecise, meaning that it is more uncertain and further away from the true value. For the parameters in the model, we obtained MCSE values between 0.00001 and 0.04177, with a mean of 0.01210 and median of 0.00659.

The code implementing the MCMC sampler, posterior summary, model diagnostics, and predictions, as well as the associated full conditional distributions of the model parameters and latent variables, are provided in the two supplemental materials as well as in the [GitHub](https://github.com/fpabonrodriguez/Research-Project-BHM) repository. In addition, this repository includes a simulation study that was conducted to support the proposed model specification.

### 3.6 Posterior Predictive Distribution and P-Values

Although the structure of a posterior predictive distribution can sometimes be calculated analytically, simulations are frequently used to obtain such forecasts. In this work, disease progression forecasts were obtained using a sampling process (simulations) from the posterior distributions, following these steps:

i. Sample a value from the posterior distributions of each parameter,
ii. Obtain the estimated model components (predicted outcomes) from the Bayesian model specification in Equation 3, and
iii. Repeat steps (i) and (ii) for *S* = 1000 times (*s* = 1, 2, 3 …, *S*), where *S* represents the number of simulations performed.

From the posterior predictive distributions, we can estimate Bayesian p-values, which are a statistical method used to assess the fit of a model to observed data^27^. They measure the probability of observing data as extreme or more extreme than the observed data, given the model and the prior distribution. In the Bayesian framework, a posterior predictive p-value is an important tool for model checking and selection^32^. It allows us to evaluate whether the model adequately explains the observed data and whether the model is consistent with prior knowledge and assumptions. This approach can help identify potential flaws in the model and guide the choice of alternative models. To compute the posterior predictive p-value, a test statistic is required to summarize the observed data^32^. This test statistic is then compared to a distribution of the same test statistic generated from samples from the posterior predictive distribution. In the current work, we used the mean squared error (MSE) as the test statistic. To obtain the posterior predictive p-values, we followed the steps:

i. Fit a Bayesian model to the data and obtain samples from the posterior distribution using MCMC methods
ii. Generate new data from the posterior predictive distribution using the sampled parameters from the posterior distribution as described at the beginning of this subsection
iii. Compute the test statistic on the observed data and on each simulated dataset from the posterior predictive distribution
iv. Calculate the posterior predictive p-value as the proportion of simulated test statistics that are as extreme or more extreme than the observed test statistic

## 4 RESULTS

The trajectories of the observed pathogen loads and antibody levels over each of the seven-time points, each separated by three months, are shown in Figure 2. The black line shows the mean trajectory of the subjects. This cohort enrolled dogs with subclinical CanL, therefore while most subject’s parasitemia and antibody level trajectories illustrate a steady state or slow progression of the disease, we also observe several examples indicative of rapid changes to the immunopathogenic state, since the loss of immune control over pathogen replication is known to occur as the disease progresses. This low production of antibodies agrees with previous findings in dogs with subclinical or early disease. Increasing antibody levels are seen in a subset of dogs, which has been shown to occur in dogs experiencing clinical progression.

**FIGURE 2.**
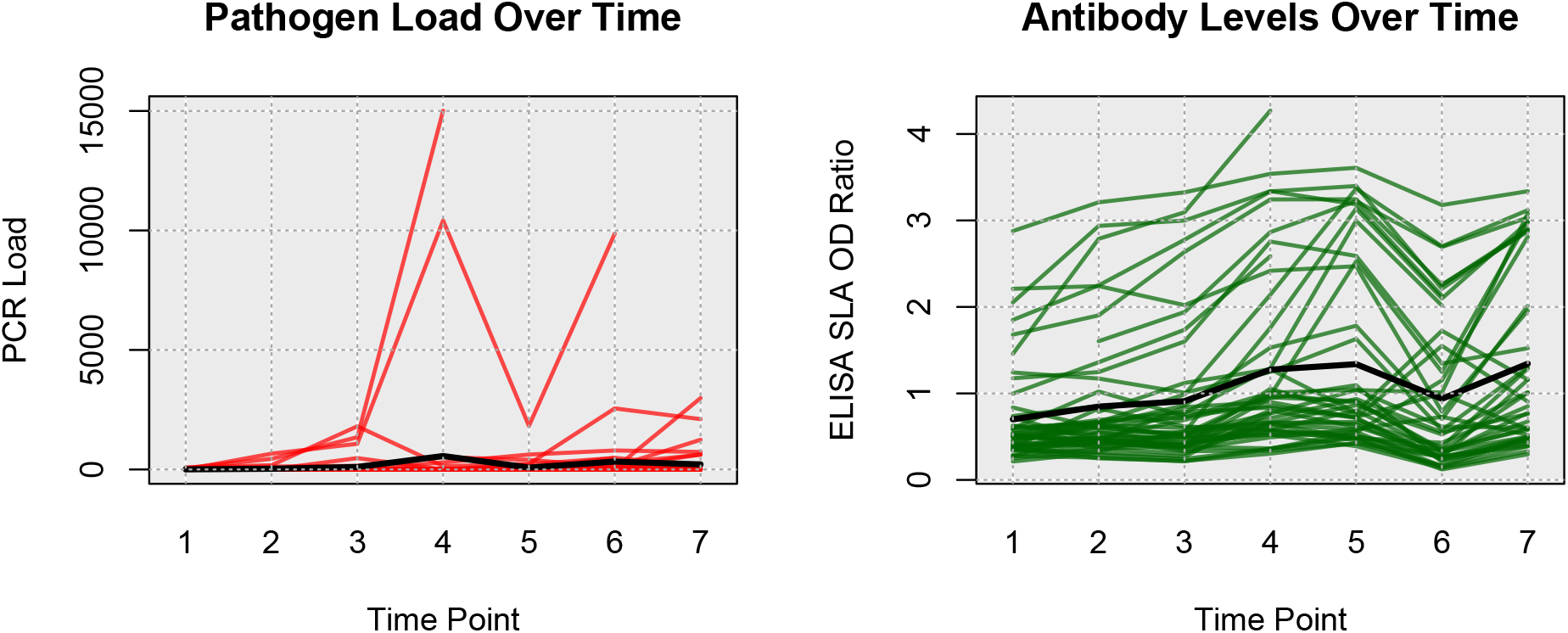
Pathogen Load and Antibody Level Kinetics: (Left plot) *Leishmania* pathogen load over time, expressed as the number of parasites per mL of blood. (Right plot) Anti-*Leishmania* antibody level over time expressed as the ratio to control cut-off. In both plots, the black line represents mean population trajectories. Each line represents a single dog’s trajectory. Each time point corresponds to 3 elapsed months.

The number of dogs classified into each disease stage over time is shown in Table 1. At the beginning of the study, most of the dogs were classified as either healthy or subclinical; the remaining two categories are clinical and removed. Due to unforeseen circumstances, the disease status for the second time point (month 6) was not able to be collected, and some other missing values were also reported in different time points for pathogen load and antibody levels due to random external events unrelated to the status of the dogs. Since these entries were not directly observed, they are considered latent variables and were imputed within the Bayesian analysis. The last two rows of the table indicate the number of unobserved entries at each time point, and the total number of subjects at each time point, which is 50, respectively.

**TABLE 1.** Number of dogs in each disease status over time. Observed number of dogs in each category of the disease state component of the Bayesian model. Post-follow-up time points after the study ended were available but not included in the current analysis.

| Disease Status | Time Point |  |  |  |  |  |  |
| --- | --- | --- | --- | --- | --- | --- | --- |
|  | 1 | 2 | 3 | 4 | 5 | 6 | 7 |
| 1 = Healthy | 32 | 0 | 26 | 27 | 23 | 21 | 19 |
| 2 = Mild | 16 | 0 | 21 | 18 | 22 | 19 | 19 |
| 3 = Severe | 2 | 0 | 0 | 1 | 0 | 4 | 2 |
| 4 = Removed | 0 | 0 | 0 | 0 | 1 | 1 | 3 |
| Missing (NA) | 0 | 50 | 3 | 4 | 4 | 5 | 7 |
| Total | 50 | 50 | 50 | 50 | 50 | 50 | 50 |

In terms of the study’s findings, we found moderate to strong evidence of temporal dependence across pathogen load, antibody levels, and illness status throughout CanL infection. This means that previous behaviors of these components had a moderate to strong influence on their future values. To summarize these effects, explanations, and interpretations of the results from this Bayesian analysis are presented, which pertains to Table 2. In general, this table shows a summary of the posterior results for all parameters in the model, including posterior means, standard deviation (SD), 95% credible intervals (Cr-I), and posterior probabilities for each parameter of being positive, which are used for assessing the strength of evidence for corresponding parameters. The posterior probabilities of positive lag-1 dependence were 0.840 for pathogen load, and approximately 1 for antibody levels, and disease status 1 and 2, respectively. In contrast, the effect for disease status 3 was negative (posterior probability of negative lag-1 is 0.874). This last result could be due to the fact that dogs in clinical stage 3 (severe) have presumably developed regulatory responses against the infection at a given time, leading to the production of important immunological cells such as CD4+ and CD8+ T cells, B cells, and macrophages^33,34,35^, which can cause a reduction and control of pathogen production.

**TABLE 2.** Posterior Summary Results. Posterior summary of the three MCMC chains. In this summary, posterior means, standard deviation, and 95% credible intervals are provided, organized by outcome model (pathogen, antibodies, and disease status). In addition, posterior probabilities of the parameters being positive are included in the last column. A brief description of these parameters is presented in the second column. The symbol * represents an interaction of the indicated model elements. The terms expressed as differences in pathogen load (or antibody levels) are denoted by 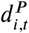 or 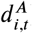 within the model specification.

| Parameter | Interpretation | Mean | SD | 95% Cr-I | Prob > 0 |
| --- | --- | --- | --- | --- | --- |
| **Pathogen Load (P)** |  |  |  |  |  |
| $\beta_{P,1}$ | Pathogen Load | 0.7049 | 0.7243 | [-0.6768, 2.2512] | 0.8433 |
| $\beta_{P,2}$ | Pathogen Load * (D=1) | 0.1707 | 0.8839 | [-1.5822, 1.8942] | 0.5767 |
| $\beta_{P,3}$ | Pathogen Load * (D=2) | 0.4538 | 0.7250 | [-1.0940, 1.8612] | 0.7506 |
| $\beta_{P,4}$ | Pathogen Load * (D=3) | -0.0323 | 0.9391 | [-1.8752, 1.7981] | 0.4868 |
| $\beta_{P,5}$ | (D=1) * Difference in Pathogen Load | 0.1183 | 0.9555 | [-1.7628, 2.0052] | 0.5483 |
| $\beta_{P,6}$ | (D=2) * Difference in Pathogen Load | -0.8736 | 0.1237 | [-1.1104, -0.6468] | 0.0006 |
| $\beta_{P,7}$ | (D=3) * Difference in Pathogen Load | -0.0102 | 0.9767 | [-1.9241, 1.8926] | 0.4942 |
| $\beta_{P,8}$ | (D=1) * Difference in Antibody Levels | 0.0163 | 0.0406 | [-0.0587, 0.0925] | 0.6698 |
| $\beta_{P,9}$ | (D=2) * Difference in Antibody Levels | -0.0030 | 0.0507 | [-0.0822, 0.0754] | 0.4644 |
| $\beta_{P,10}$ | (D=3) * Difference in Antibody Levels | 0.0525 | 0.2307 | [-0.3916, 0.5111] | 0.5929 |
| $\sigma_P^2$ | Variance in Pathogen Load | 0.0215 | 0.0058 | [0.0194, 0.0231] | - |
| $\alpha_{P,1}$ | Intercept | -0.0005 | 0.0095 | [-0.0160, 0.0144] | 0.4813 |
| $\alpha_{P,2}$ | Age group | 0.0017 | 0.0056 | [-0.0025, 0.0063] | 0.8276 |
| $\alpha_{P,3}$ | SNAP results | 0.0020 | 0.0044 | [-0.0037, 0.0074] | 0.7409 |
| $\alpha_{P,4}$ | DPP results | 0.0103 | 0.0100 | [-0.0041, 0.0250] | 0.9211 |
| $\alpha_{P,5}$ | Treatment group | -0.0027 | 0.0050 | [-0.0082, 0.0026] | 0.1334 |
| **Antibodies Level (A)** |  |  |  |  |  |
| $\beta_{A,1}$ | Pathogen Load | -0.1662 | 0.3397 | [-0.7823, 0.4588] | 0.2894 |
| $\beta_{A,2}$ | Antibody Levels | 0.8976 | 0.0589 | [0.7931, 1.0084] | 0.9999 |
| $\beta_{A,3}$ | (D=1) * Difference in Pathogen Load | 0.6141 | 0.9863 | [-1.3045, 2.5770] | 0.7330 |
| $\beta_{A,4}$ | (D=2) * Difference in Pathogen Load | 0.4121 | 0.2532 | [-0.0764, 0.8862] | 0.9510 |
| $\beta_{A,5}$ | (D=3) * Difference in Pathogen Load | 0.0888 | 0.9938 | [-1.8698, 2.0246] | 0.5364 |
| $\beta_{A,6}$ | (D=1) * Difference in Antibody Levels | -0.3924 | 0.1008 | [-0.5912, -0.1977] | 0.0003 |
| $\beta_{A,7}$ | (D=2) * Difference in Antibody Levels | -0.1200 | 0.1137 | [-0.3297, 0.0877] | 0.1280 |
| $\beta_{A,8}$ | (D=3) * Difference in Antibody Levels | -0.1632 | 0.4645 | [-1.1384, 0.7042] | 0.3730 |
| $\sigma_A^2$ | Variance in Antibody Levels | 0.0515 | 0.0039 | [0.0473, 0.0560] | - |
| $\alpha_{A,1}$ | Intercept | -0.0079 | 0.0179 | [-0.0433, 0.0279] | 0.3129 |
| $\alpha_{A,2}$ | Age group | -0.0034 | 0.0079 | [-0.0137, 0.0076] | 0.2705 |
| $\alpha_{A,3}$ | SNAP results | 0.0141 | 0.0067 | [0.0009, 0.0267] | 0.9811 |
| $\alpha_{A,4}$ | DPP results | 0.0470 | 0.0229 | [-0.0004, 0.0876] | 0.9747 |
| $\alpha_{A,5}$ | Treatment group | 0.0048 | 0.0077 | [-0.0081, 0.0172] | 0.7721 |
| **Disease Status (D)** |  |  |  |  |  |
| $\beta_{D1,1}$ | (D=1) | 1.8619 | 0.2182 | [1.4430, 2.2855] | 0.9999 |
| $\beta_{D1,2}$ | (D=1) * Pathogen Load | 0.0064 | 0.9901 | [-1.9069, 1.9523] | 0.5044 |
| $\beta_{D1,3}$ | (D=1) * Antibody Levels | -0.1016 | 0.9536 | [-1.9728, 1.7355] | 0.4633 |
| $\beta_{D2,1}$ | (D=2) | 1.7409 | 0.2527 | [1.2549, 2.2433] | 1 |
| $\beta_{D2,2}$ | (D=2) * Pathogen Load | 0.0469 | 0.9886 | [-1.8529, 1.9786] | 0.5105 |
| $\beta_{D2,3}$ | (D=2) * Antibody Levels | 0.3826 | 0.9035 | [-1.4137, 2.1628] | 0.6617 |
| $\beta_{D3,1}$ | (D=3) | -0.7736 | 0.6895 | [-2.1861, 0.5285] | 0.1263 |
| $\beta_{D3,2}$ | (D=3) * Pathogen Load | -0.0041 | 0.9991 | [-1.9845, 1.9494] | 0.4983 |
| $\beta_{D3,3}$ | (D=3) * Antibody Levels | 0.1259 | 0.9875 | [-1.8258, 2.0582] | 0.5521 |
| $\alpha_{D,1}$ | Intercept | 0.6887 | 0.9354 | [-1.1200, 2.5174] | 0.7679 |
| $\alpha_{D,2}$ | Age group | 0.9730 | 0.8063 | [-0.5359, 2.6314] | 0.8853 |
| $\alpha_{D,3}$ | SNAP results | 0.9014 | 0.8253 | [-0.6646, 2.5676] | 0.8610 |
| $\alpha_{D,4}$ | DPP results | 0.0473 | 0.9994 | [-1.8891, 2.0141] | 0.5183 |
| $\alpha_{D,5}$ | Treatment group | 0.9217 | 0.8372 | [-0.6550, 2.6156] | 0.8682 |

We found that the dependence of pathogen load on previous values varied by stage; the conditional interaction effect on pathogen load of lag-1 change in pathogen load for stage 2 dogs had a posterior mean of −0.874 with 95% Cr-I [−1.110, −0.647] and posterior probability 0.999 that the true parameter is less than zero. This result suggests that subjects in the mild stage of the disease could show some decrease in the pathogen burden as time passes, which is consistent with the discussion from Petersen et al.^36^, where it was shown that parasite load tends to decrease once certain inflammatory responses from the immune system are active. We find moderate evidence that co-infection status (measured via 4Dx SNAP test) and dog age were associated with increased parasite load overall (posterior probabilities of being positive are 0.740 and 0.828), as well as that the sarolaner treatment group had lower overall parasite loads (posterior probability of being negative of 0.860). Consistent with expectation, we also find compelling evidence that stronger DPP responses were associated with higher pathogen burden (posterior probability of being positive of 0.921).

In terms of antibodies, several compelling effects emerge. First, we find strong evidence that co-infection status (4Dx SNAP test) was associated with increased anti-Leishmania antibody levels, with a posterior probability of being positive of 0.981. This effect has a posterior mean of 0.014 with 95% Cr-I [0.001, 0.027]. Testing for tick-borne bacterial antigens can help us understand how the presence of other pathogens may affect the production of anti-Leishmania antibody levels and the general inflammatory response^18^. In addition, stronger DPP responses were associated with higher antibody levels (posterior probability 0.975), which is expected since the DPP serological test also detects anti-*Leishmania* antibodies similar to SLA ELISA. We also found that the conditional interaction effect on antibody levels of lag-1 change in anti-Leishmania antibody levels for stage 1 dogs had a posterior mean of −0.392 with 95% Cr-I [−0.591, −0.198] and posterior probability approximately 1 that the true parameter is less than zero.

In addition to characterizing evidence for the processes driving disease progression, we also investigated the degree to which the presented Bayesian model can predict individual and aggregate patterns of CanL progression. Using posterior predictive distributions based on parameters drawn from the converged MCMC samples, we investigated different outcome patterns. For this context, we identified four dogs within the same cohort based on four different outcomes scenarios, from decreasing or stable condition to growing clinical status. Subject 1, which is considered under the rapid progression scenario, goes from a subclinical state to a severe condition. It was seropositive at a low level at enrollment and showed a coinfection with *Ehrlichia spp*. It experienced a tick-borne coinfection at timepoint (TP) 3. The subject experienced a rapid increase in antibody levels and parasitemia across each TP1 to TP4. The subject jumped from mild disease at TP3 to severe clinical presentation at TP4 and was euthanized for CanL at TP4. Subject 2, which is a non-progressor, was seronegative at enrollment and did not experience a tick-borne coinfection. The subject remained seronegative, PCR negative, and subclinical throughout the study only varying from LeishVet stage 0 to 1. Subject 3 was considered a gradual progressor. It was seronegative at enrollment. Between TP3 and TP4, it tested positive for *Borrelia burgdorferi* and *Anaplasma spp*. It remained *B. burgdorferi* seropositive for the duration of the study. At TP4, the subject showed borderline parasitemia that progressed to positive for the duration of the study. LeishVet score also progressed from level 2 to level 3 (mild to moderate disease) between TP5 and TP6. The last subject considered, Subject 4, was under the stable scenario. It was seronegative at enrollment and never tested positive for tick-borne coinfection. The subject became relatively low, stably seropositive at time points 5 to 7 followed by low, stable parasitemia. The clinical score remained stable at LeishVet stage 1 and showed some improvement to stage 0 at the final time point. The goal was to predict the pathogen load, antibody level, and probabilities of the dogs to be classified in each disease status for additional points in time, assuming that only the information from the first two time points was available, and compare estimated trajectories from the available observed trajectories.

For pathogen load and antibody level, we plot the predicted trajectories over time for each of the subjects, as well as the marginal probabilities of being in a particular disease state at each time point. We hypothesize that disease patterns can be predicted using diagnostic parameters with 6 months between measurements, as suggested by veterinarians. Assessing certain diagnostic variables every 6 months allows veterinarians to monitor a dog’s health over time and detect changes that may indicate the onset or progression of the disease. Our model was constructed in such a way that allows us to use available data from the immediate past or present, and predict outcomes farther away in time. More frequent follow-up appointments may not be feasible or necessary for many dogs, while longer intervals between appointments may allow changes in health status to go undetected for too long. Since predictions in a 6 months interval were preferred, we can still use the information on a 3-month basis and make the predictions. Clearly, the amount of available data and the frequency of observed progression events will affect the predictive performance of the model.

Figures 3 and 4 show the predicted pathogen load and antibody level trajectories for the four subjects, based on different outcome patterns, respectively. For each prediction, we present 80% and 95% prediction intervals, and a line representing the mean trajectories. In addition, we plotted the observed trajectories of the dogs. From this simple prediction setting, we were able to capture some of the observed trajectories of the dogs for the antibody levels. For pathogen load, the mean trajectory of the simulations was higher than the observed data for 3 out of these 4 dogs, and close to the observed data for the first dog. A reason for this could be that Subject 1 had a progressed disease and more completed observations. The posterior predictive p-values for pathogen load and antibody levels readouts are 0.701 and 0.766, respectively, based on the four subjects that were considered within the scenarios. These posterior p-values indicate that the observed data are somewhat consistent with the model and prior assumptions, but there is still a relatively high level of unexpectedness in the data.

**FIGURE 3.**
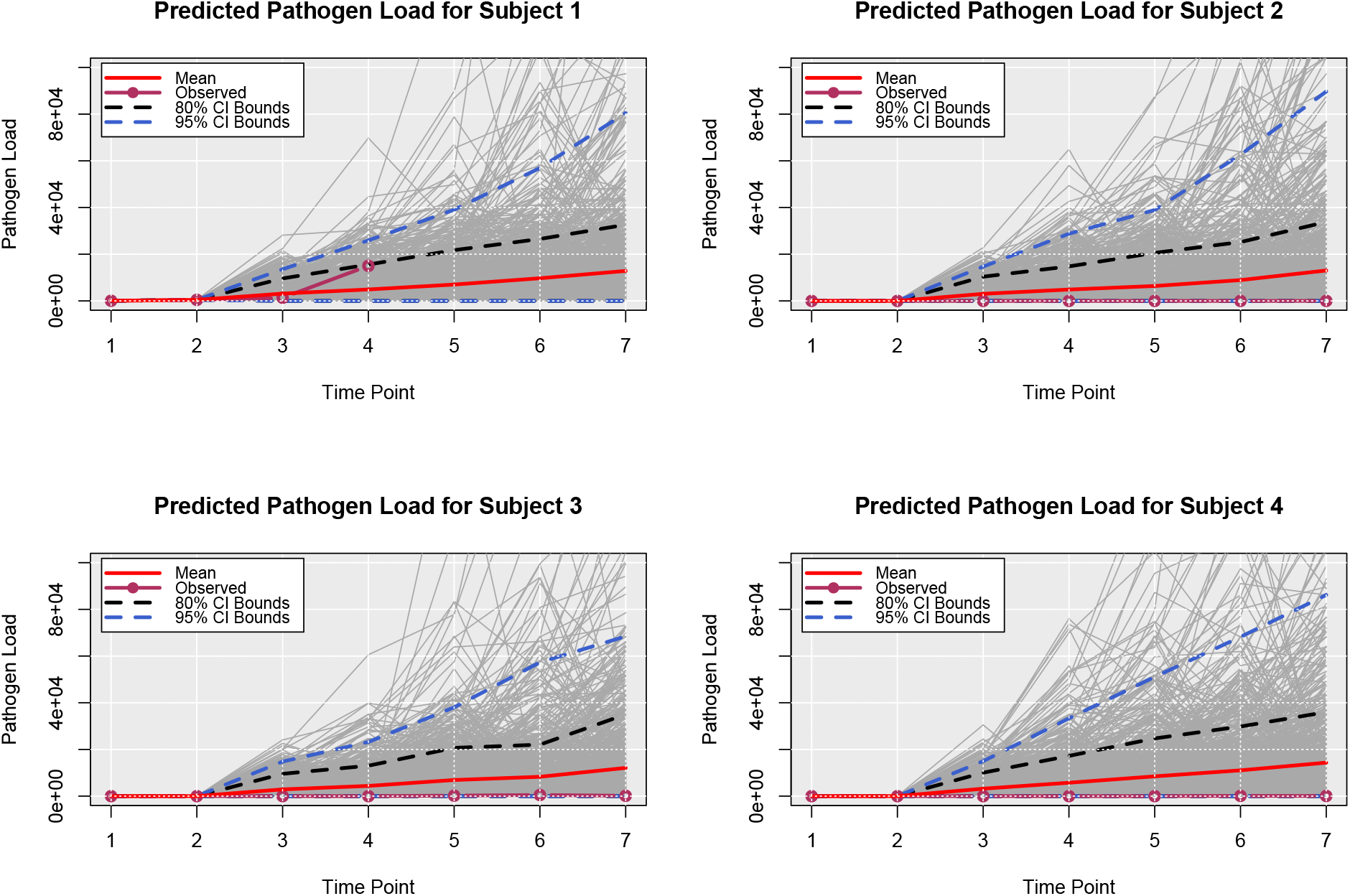
Posterior Predictive Trajectories of Pathogen Load. Posterior predicted pathogen load for four dogs. Time points 3 through 7 were predicted times.

**FIGURE 4.**
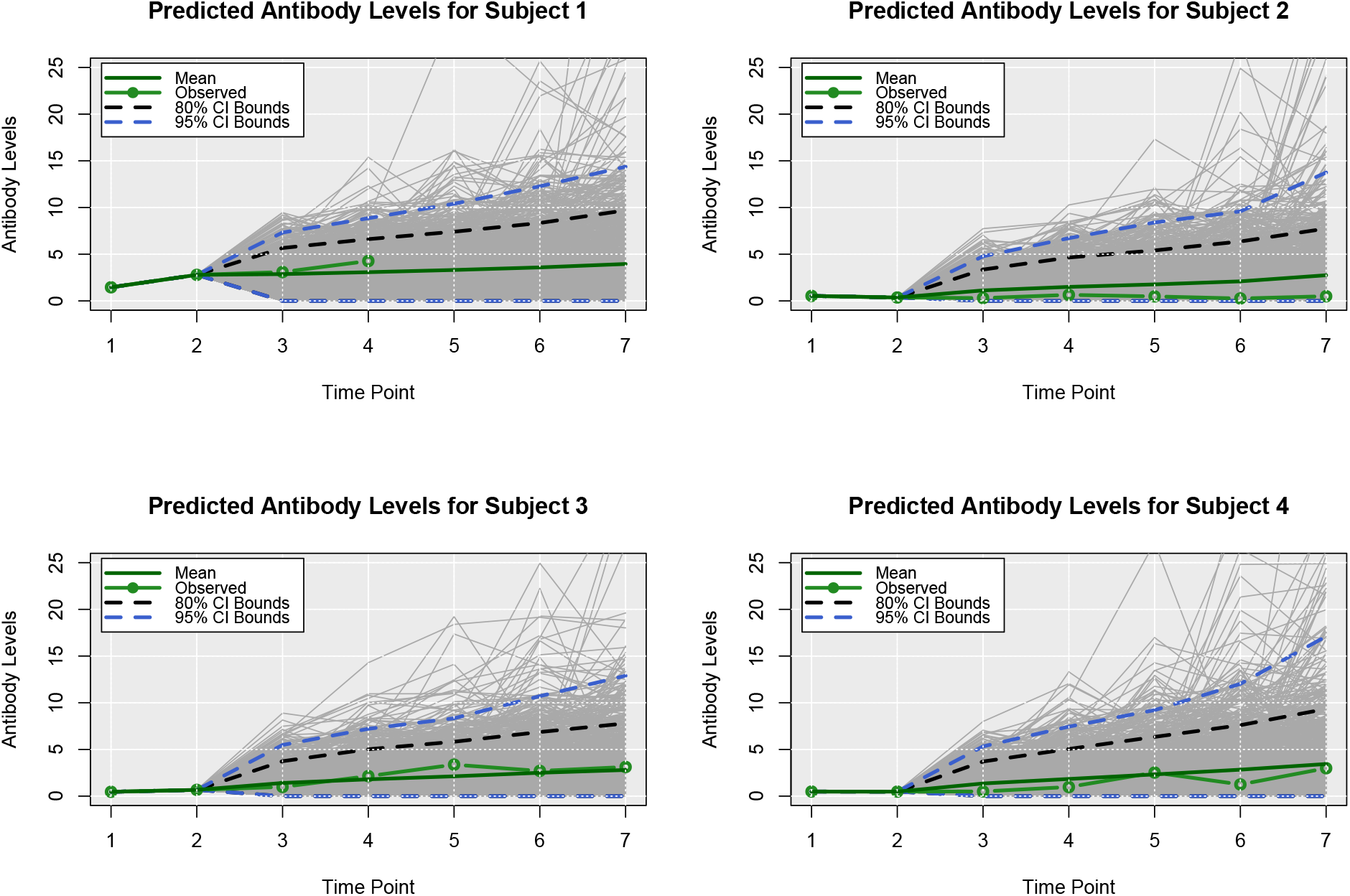
Posterior Predictive Trajectories of Antibody Levels. Posterior predicted antibodies level for four dogs. Time points 3 through 7 were predicted times.

In Figure 5, we present the posterior predicted marginal probabilities of the dogs being classified in a particular disease state over time. From these plots, we observe that the probability of being removed increases from 0% to approximately 15% for all of hese dogs, no matter their initial status. In addition, the probability of staying healthy reduced to approximately 38% for dogs 2 and 4, but it increases to around 35% for dogs 1 and 3. The presented model was able to capture several expected behaviors that were backed up by research. Dogs treated with sarolaner, for example, had lower total parasite burdens, and DPP serological test findings were linked to a higher pathogen burden. Our approach will benefit clinicians and veterinarians by allowing them to better understand immune responses and *Leishmania* infection control throughout time.

**FIGURE 5.**
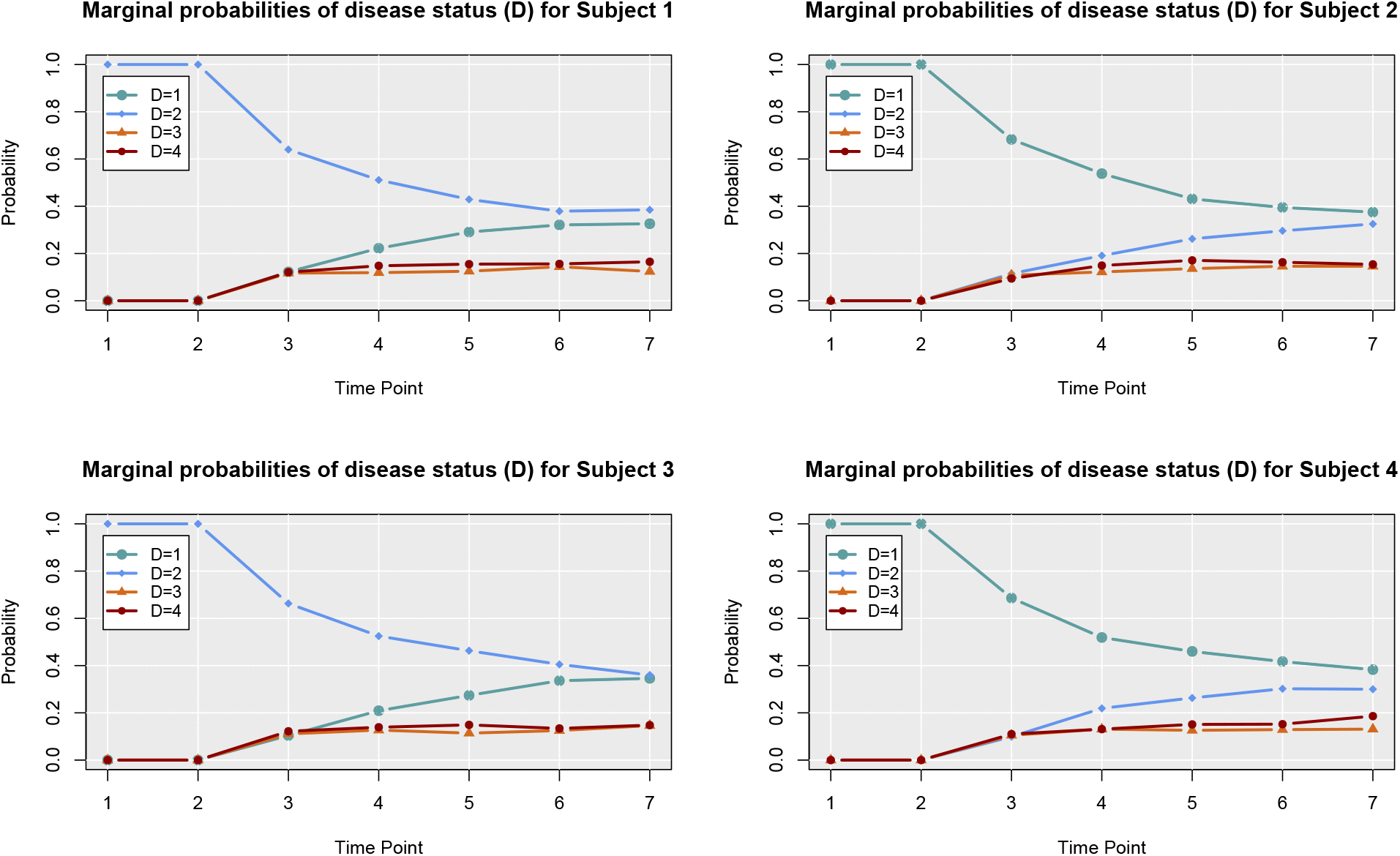
Posterior Marginal Probabilities of Disease Status. Posterior predicted (marginal) probabilities of four dogs being in a particular disease state over time. Time points 3 through 7 were predicted times. Timepoint 1 corresponds to the observed disease state and MCMC estimated latent values were used for time point 2.

## 5 DISCUSSION

There are several competing statistical and mathematical modeling approaches that could be applied to this problem, and to the best of our knowledge there is no single traditional approach that has universal acceptance in the proposed modeling setting. In the case of Frequentist methods, we felt that relying on asymptotic arguments for the short longitudinal follow-up would be sub-optimal. Moreover, we have a strong philosophical preference for Bayesian models, which are less likely to make strong and incorrect conclusions regarding hypotheses of interest due to the shrinkage effect of prior distributions^32^ and have a strong axiomatic foundation^37^. When considering mathematical models, we felt that Bayesian methods are preferable where practical due to their ability to formally quantify uncertainty concerning model parameters, rather than relying on simple optimization to provide a “best fit”. From the same Figure 1, we saw that at each time point, the observed and latent measures which characterize the pathogen-host dynamics and their interactions comprise the state of the model. Using a conditional structure under distributional assumptions is one way to model the evolution of a disease^32,27^, allowing us to draw conclusions and quantify uncertainty about important variables related to the immune response to infections. In addition, the Bayesian setting also enables us to easily handle unobserved values as latent quantities and to estimate them from the available data and model structure^26^.

Through this statistical analysis, we were able to explore and characterize some evidence for the processes that drive CanL progression and the host immune response to infection. We investigated the degree to which the presented Bayesian model can predict individual and aggregate patterns of disease progression. We noticed that even when the observed data was collected in 3-month intervals, making predictions in this time frame is not recommended since the disease progression pattern may not yet be apparent. As shown with the forecasting scenarios, the presented model specification is still considered to be too simple to fully understand the disease progression and immune responses, and further considerations need to be undertaken to model this complex disease. For instance, considering CD4+ T proliferation over time, and information on different important cytokine expressions such as protective interferon-gamma (IFN-gamma) or inhibitory interleukin 10 (IL-10) could improve the model and its predictions^3^. Although the induction of CD4+ T helper 1 cell responses is considered essential for immunity against *Leishmania*, B cells and the production of *Leishmania*-specific antibodies have also been proposed to play an important role in disease progression^18^.

While posterior predictive plots and leave-one-out cross-validation (LOOCV) are both techniques used in Bayesian data analysis to evaluate the fit of a statistical model, in this work we relied on the former technique. Posterior predictive plots involve simulating data from the fitted model and comparing it to the observed data. This allows us to visually assess how well the model can reproduce the patterns and variability in the observed data. On the other hand, LOOCV involves fitting the model to all but one of the data points and then using the model to predict the value of the omitted point. This is repeated for each data point, and the predicted values are compared to the observed values. This allows us to evaluate how well the model can predict new data based on the information in the existing data. A future direction of this work will use a combination of techniques, including posterior predictive plots, LOOCV, and other model diagnostic tests, to fully evaluate the fit of our proposed modeling approach.

An extended version of this model will consider additional important immunological parameters controlling CanL disease presentation, such as *Leishmania*-antigen-specific CD4+ T cell proliferation, IL-10, and IFN-gamma production. Based on Nylén and Gautam^38^, high levels of *Leishmania* specific antibodies are observed in subjects with *Leishmania* infection and other severe forms of Leishmanial disease and there is accumulating evidence that B cells and antibodies correlate with pathology. Future work incorporating these enhancements, and additional post-study longitudinal follow-up of the canine cohort should enable us to better understand CanL progression and to refine the predictive performance of the models to create a useful clinical tool.

## 6 ACKNOWLEDGEMENTS

Research reported in this article was supported by the National Institutes of Allergy and Infectious Diseases (NIAID) of the National Institutes of Health (NIH) of the United States of America under award number R01AI139267-03 and by MFHA18441000. This work was also performed while B.M.S. was supported by NIH/NIAID T32AI007260, which was the basis for data collection. The content is solely the responsibility of the authors and does not necessarily represent the official views of the National Institutes of Health.

## 8 DATA AVAILABILITY STATEMENT

The authors confirm that the data analyzed supporting the findings of this study are available within the article and supplementary materials.

